# Multiple competing RNA structures dynamically control alternative splicing in human ATE1 gene

**DOI:** 10.1101/2020.06.04.134791

**Authors:** Marina Kalinina, Dmitry Skvortsov, Svetlana Kalmykova, Timofei Ivanov, Olga Dontsova, Dmitri D. Pervouchine

## Abstract

The mammalian *Ate1* gene encodes an arginyl transferase enzyme, which is essential for embryogenesis, male meiosis, and regulation of the cytoskeleton. Reduced levels of *Ate1* are associated with malignant transformations and serve as a prognostic indicator of prostate cancer metastasis. The tumor suppressor function of *Ate1* depends on the inclusion of one of the two mutually exclusive exons (MXE), exons 7a and 7b. Here, we report that the molecular mechanism underlying MXE splicing in Ate1 involves five conserved regulatory intronic elements R1–R5, of which R1 and R4 compete for base pairing with R3, while R2 and R5 form an ultra-long-range RNA structure spanning 30 Kb. In minigenes, single and double mutations that disrupt base pairings in R1R3 and R3R4 lead to the loss of MXE splicing, while compensatory triple mutations that restore the RNA structure also revert splicing to that of the wild type. Blocking the competing base pairings by locked nucleic acid (LNA)/DNA mixmers complementary to R3 leads to the loss of MXE splicing, while the disruption of the ultra-long-range R2R5 interaction changes the ratio of mutually exclusive isoforms in the endogenous *Ate1* pre-mRNA. The upstream exon 7a becomes more included than the downstream exon 7b in response to RNA Pol II slowdown, however it fails to do so when the ultra-long-range R2R5 interaction is disrupted. In sum, we demonstrated that mutually exclusive splicing in *Ate1* is controlled by two independent, dynamically interacting and functionally distinct RNA structure modules. The molecular mechanism proposed here opens new horizons for the development of therapeutic solutions, including antisense correction of splicing.

## Introduction

Arginylation is a widespread post-translational protein modification that transfers an L-arginyl residue from the Arg-tRNA onto the polypeptide chain (*1*). It is mediated by the arginyl transferase encoded within *Ate1* gene (*2*). *Ate1* is essential in most eukaryotic systems and is implicated in regulation of many physiological pathways including proteolysis (*3,4*), response to stress and heat shock (*5–7*), embryogenesis (*8–10*), regenerative processes (*11–13*), and aging (*14, 15*). *Ate1* has recently been identified as a master regulator affecting disease-associated pathways (*16–18*), and its knockout results in embryonic lethality and severe developmental defects in mice (*9,10,19,20*).

Like many other eukaryotic genes, *Ate1* generates several mRNA isoforms through alternative splicing (*21*). In mammals, they differ by mutually exclusive choice between two adjacent, homologous 129-bp exons (7a or 7b), and by alternative choice of the initial exon (1a or 1b) (*22*). The two major mRNA isoforms of Ate1 are *Ate1-1* (1b7a) and *Ate1-2* (1b7b), and the isoforms that contain both exon 7a and 7b are suppressed by mutually exclusive splicing (*18*). In mice, *Ate1-1* and *Ate1-2* are expressed stably in all tissues, but their ratio varies from 0.1 in the skeletal muscle to 10 in the testis (*21,23,24*). While *Ate1-2* is almost completely cytosolic, *Ate1-1* localizes in both cytosol and nucleus (*21*) and can specifically interact with *Liat1*, a testis-specific molecule (*25*). Furthermore, Ate1-knockout cells can form tumors in subcutaneous murine xenograft assays, in which the tumor growth can be partially rescued by the reintroduction of stably expressed *Ate1-1*, but not *Ate1-2* (*16*). The isoforms *Ate1-3* (1a7a) and *Ate1-4* (1a7b) encode a variant of arginyl transferase that is specific for N-terminal cysteine, with tissue-specific expression, cellular localization, and carcinogenic potential similar to those of *Ate1-1* and *Ate1-2*, respectively (*22*). The ratio of *Ate1* isoforms containing exons 7a and 7b switches substantially during male meiosis in mice suggesting a role in the mitotic-to-meiotic transition of the germ cell cycle (*24*). All these observations suggest that the sequences of amino acids encoded by exons 7a and 7b result in functionally distinct arginyl transferases.

To date, the mechanism of MXE splicing of exons 7a and 7b in *Ate1* has not been studied in detail (*18*). Generally, MXE clusters are believed to evolve through tandem genomic duplications (*26–28*), and many of them implement mutually exclusive splicing through a mechanism that depends on competing RNA structures (*29*). Transcripts of such genes, of which *Dscam* (*30*) is the most prominent example, contain multiple sites called the selector sequences that are complementary to a regulatory element called the docking site. Only one of the competing base pairings can form at a time, which exposes only one exon in a cluster to the spliceosome and eventually leads to mutually exclusive splicing (*29*). In complex cases, several groups of docking and selector sequences can operate on a single gene creating a bidirectional control mechanism (*31*). A recent genome-wide study of MXE clusters conjectured that docking/selector systems may evolve spontaneously as a natural byproduct of tandem duplications, in which only one of the two arms of an ancestral stem-loop is duplicated (*32, 33*).

Since exons 7a and 7b of *Ate1* are homologous (nucleotide sequence identity 45%) and similar in length, most likely having originated through tandem genomic duplications, we questioned here whether their splicing is also controlled by competing RNA structures. Using comparative sequence analysis, we identified five regulatory intronic elements in *Ate1* pre-mRNA and examined their function in alternative splicing by means of site-directed mutagenesis and locked nucleic acid (LNA)/DNA mixmers as antisense oligonucleotides (AONs) that interfere with RNA secondary structure.

## Results

### Conserved complementary regions in exon 7 cluster

In order to identify the mechanisms responsible for mutual exclusive splicing of exons 7a and 7b, we used comparative sequence analysis to search for potential regulatory sequences in their intervening and flanking introns (Figure 1A). We found that the regions immediately upstream of exons 7a and 7b, termed here as R1 and R4, are highly similar to each other and show remarkable sequence conservation across vertebrate species. The intron between exons 7a and 7b contains two conserved regions, termed here as R2 and R3, where R3 is complementary to both R1 and R4, while R2 is complementary to another highly conserved region R5 located ~30 kb downstream in the intron flanking exon 7b. The pattern of complementarity between these regions suggests that R1 and R4 could compete with each other for base pairing with R3 and, together with base pairing of R2 with R5, they form a pseudoknot (Figure 1B).

**Figure 1:**
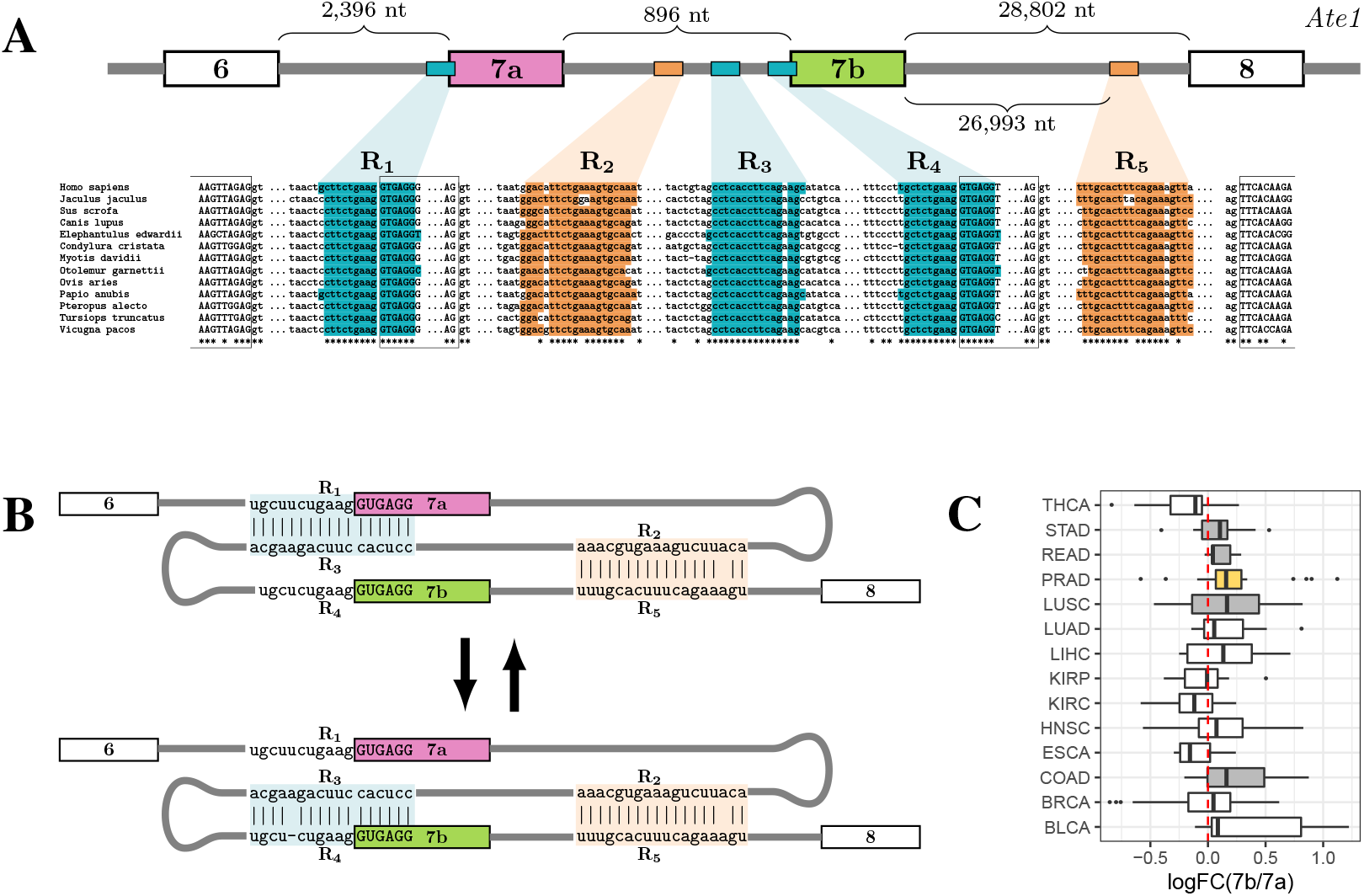
**(A)** *Ate1* contains five regulatory intronic elements (R1–R5) that are conserved through the evolution of vertebrates. **(B)** The sequences of R1 and R4 are highly similar to each other and both are complementary to R3; R2 is complementary to R5 despite 30 Kb distance. R1 and R4 can compete for base pairing with R3, and together with the R2R5 they form a pseudoknot. **(C)** The log fold change of 7b/7a isoforms (the difference of log_10_(*7b/7a*) between cancer and normal tissue) significantly increases in stomach (STAD), rectum (READ), colon (COAD), and prostate (PRAD) adenocarcinomas, and in lung squamous cell carcinoma (LUSC); Wilcoxon test FWER < 0.05 (yellow), *P* < 0.05 (gray).

To test whether exons 7a and 7b are indeed spliced in a mutually exclusive manner, we analyzed a large compendium of RNA-seq experiments from the Genotype Tissue Expression (GTEx) project (*34*) (Figure S1A). Consistent with previous reports (*17*), exons 7a and 7b have a broad range of inclusion levels across tissues (median 33% and 67%, respectively) with the most remarkable deviation in testis (median 60% and 39%, respectively). There was almost no evidence of simultaneous inclusion or simultaneous skip of both exons neither in normal tissues (Figure S1A), nor in cancer samples from The Cancer Genome Atlas (Figure S1B) (*35*). However, in agreement with previous studies (*16*), we detected a significant prevalence of isoforms containing exon 7b as compared to exon 7a in matched samples of prostate adenocarcinomas and normal tissues (FWER<0.05) and also in other epithelial tumors including stomach, rectum, colon, and lung squamous cell carcinoma (Figure 1C).

The occurrence of competing RNA structures in mutually exclusive exons in *Ate1* is reminiscent of splicing control mechanisms in other genes (*29*). We therefore analyzed in detail the function of these complementary regions using site-directed mutagenesis and antisense oligonucleotides.

### Competition between R1R3 and R3R4 controls mutually exclusive splicing

To check whether RNA structure is implicated in the regulation of splicing in *Ate1*, we created a minigene construct containing its fragment spanning between exons 6 and 8 under the constitutive CMV promoter, and quantitatively assessed splice isoforms in transfection experiments using human adenocarcinoma A549 cells (Figure 2A). In the minigene, the endogenous intron downstream of exon 7b was reduced in size to ~2 kb due to obvious limitations of cloning large fragments.

**Figure 2:**
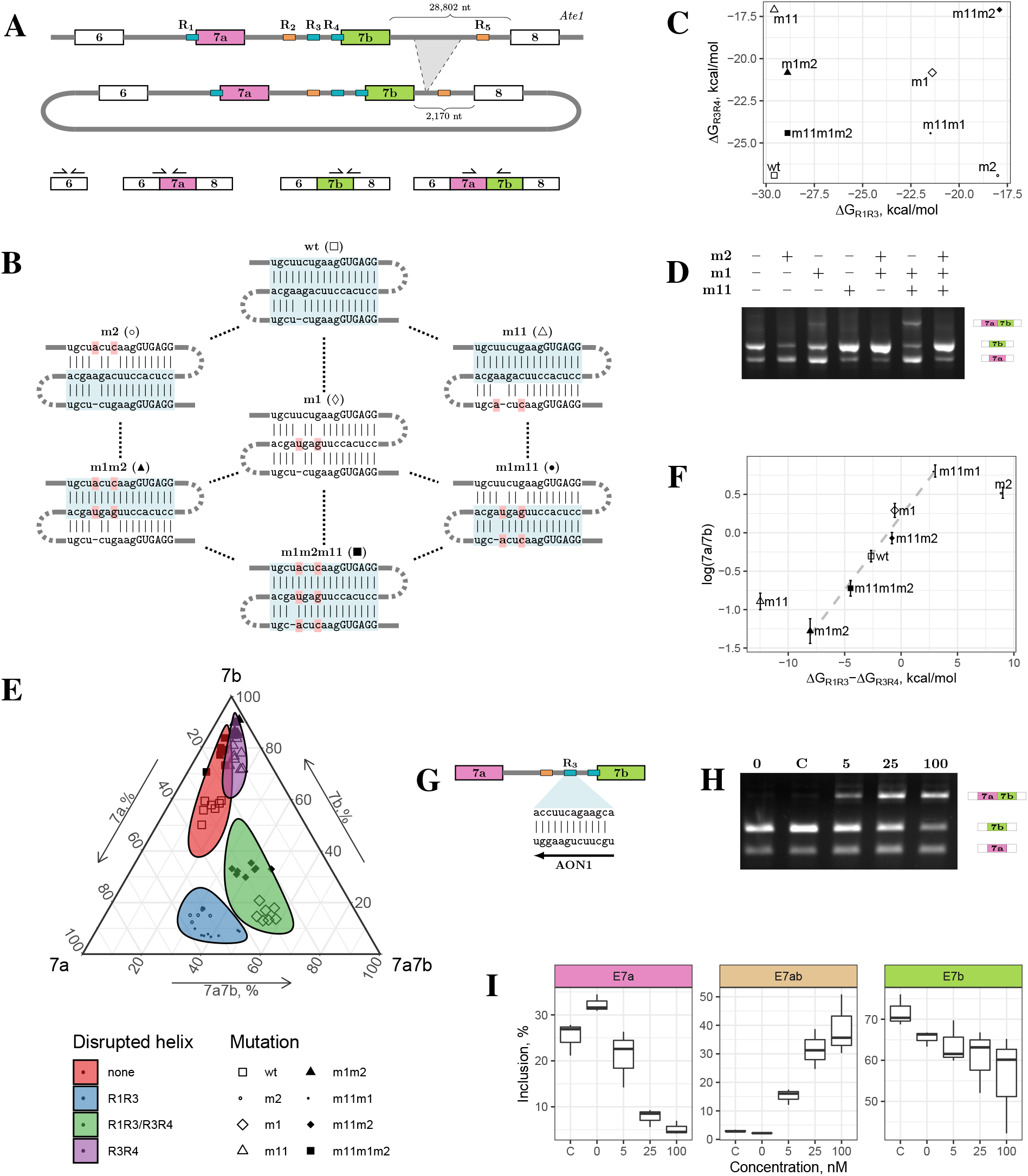
**(A)** The intron between exon 7b and exon 8 was reduced in size from ~29 Kb to ~2 Kb in the minigene (top); qRT-PCR primers for quantitative assessment of alternative splicing (bottom). **(B)** Disruptive and compensatory mutations (pink) in R1, R3, and R4. m11m2 mutant is not shown. **(C)** The predicted hybridization energies of R1R3 and R3R4 for disruptive and compensatory mutations. **(D)** Exon inclusion changes in single and double mutants; WT splicing is restored in the m1m2m11 triple mutant. **(E)** Ternary plot of exon 7a, 7b, and 7a7b inclusion. Colored areas represent 95% confidence intervals calculated via the Mahalnobis distance using log-ratio transformation (*36,37*). **(F)** The isoform ratio of exons 7a and 7b depends on the difference of thermodynamic stabilities of R1R3 and R3R4. **(G)** AON1 disrupts R1R3 and R3R4 via base pairing to R3. **(H), (I)** The inclusion of double exons increases with increasing AON1 concentration in the endogenous *Ate1* transcript. ‘C’ denotes the control AON. The scales of y-axes are different between panels.

Our strategy was to make point mutations that disrupt RNA structure when introduced alone and restore it when introduced in different combinations. First, we tested the effect of disruptive and compensatory mutations on double-stranded structures of R1R3 and R3R4 (Figure 2B and 2C). The mutation disrupting R1R3 base pairing (m2) increases the usage of exon 7a, whereas the mutation disrupting R3R4 (m11) increases the usage of exon 7b (Figure 2D and 2E). The mutated R3 (m1), which is unable to base-pair with R1 or R4, increases the proportion of transcripts containing both exon 7a and 7b. The compensatory double mutation m1m2, which restores R1R3 but disrupts R3R4, increases the efficiency of exon 7b inclusion up to 85%, which is higher than that in the wild type (WT), while exon 7a inclusion becomes almost fully suppressed. Conversely, the compensatory double mutation m1m11, which restores R3R4 while disrupting R1R3, leads to a more efficient inclusion of exon 7a compared to the WT. Finally, the proportion of splice isoforms in the triple mutant (m1m2m11), in which both R1R3 and R3R4 are restored, is more similar to that in the WT than it was in all other mutants. At that, the ratio of exon 7a/7b inclusion changed proportionally to the difference of thermodynamic stabilities of R1R3 and R3R4 in all mutants except m2 and m11 (Figure 2F). Notably, the proportion of isoforms containing both exons 7a and 7b increased whenever R1R3 base pairing was disrupted.

The cloned fragment of *Ate1* lacks a substantial part of intron 7, which may affect splicing. We therefore independently examined the role of R1, R3, and R4 in the endogenous *Ate1* transcript using locked nucleic acid (LNA)/DNA mixmers as antisense oligonucleotides (AONs) that interfere with RNA secondary structure (*38*). Since R1 and R4 overlap with splice sites, we chose to use AON with specific sequence complementary to R3 (AON1) (Figure 2G). RT-PCR analysis revealed that the transfection of 5 nM or a higher concentration of AON1 efficiently induced the inclusion of both exons 7a and 7b and suppressed the inclusion of individual exons, while the transfection of the control AON didn’t show any difference from the non-treated cells (Figure 2H). Quantitative RT-PCR (qRT-PCR) with isoform-specific primers confirmed consistent dose dependence of AON1 treatment (Figure 2I). Taken together, the mutagenesis and AON1 treatment indicate that the function of competitive base pairings between R1, R3, and R4 is to control mutually exclusive splicing of exons 7a and 7b.

### The ultra-long-range R2R5 base pairing controls isoform bias

To elucidate the function of the other two conserved regions, R2 and R5, we applied a similar mutation strategy to the minigene carrying a reduced *Ate1* fragment (Figure 3A). When introduced separately, the disruptive mutations m3 and m4 almost completely abrogated the inclusion of exon 7a and strongly enhanced the inclusion of exon 7b, while the compensatory double mutant m3m4 reverted the splicing pattern to that of the WT (Figure 3B). Remarkably, the disruptive mutations affected only the ratio of isoforms carrying mutually exclusive exons, but not the proportion of transcripts with double exons (Figure 3C).

**Figure 3:**
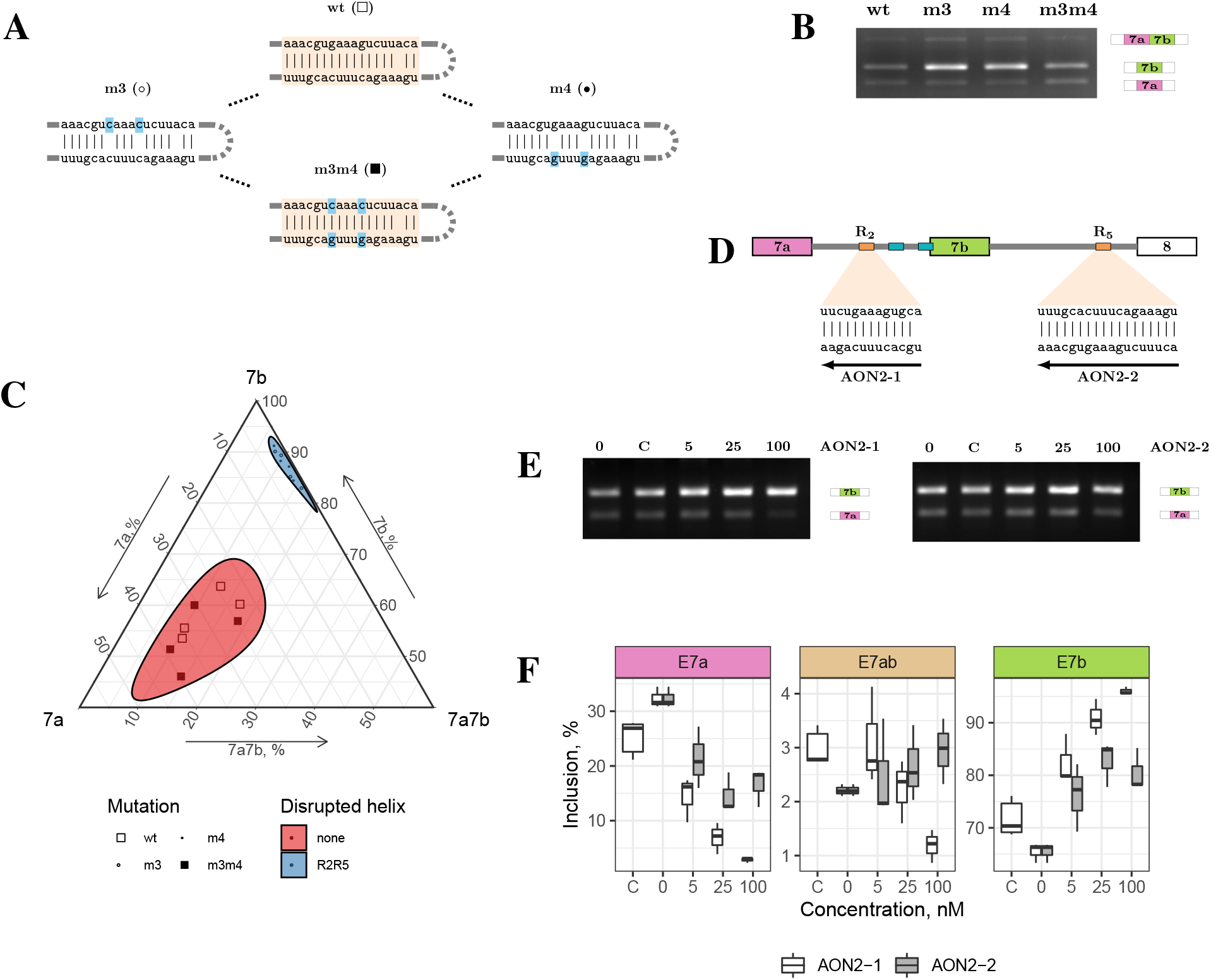
**(A)** Disruptive and compensatory mutations in R2 and R5. **(B), (C)** The mutations disrupting R2R5 base pairing (m3 and m4) promote exon 7b inclusion, while the compensatory mutation (m3m4) restores WT splicing. Ternary plots are as in Figure 2E. **(D)** Base pairing with AON2-1 and AON2-2 disrupts R2R5 interaction. **(E), (F)** Increasing concentration of AON2-1 and AON2-2 suppresses exon 7a and promote exon 7b usage without inducing double (7a7b) exons in the endogenous *Ate1* transcript. ‘C denotes the control AON. The scales of y-axes in (F) are different between panels.

In the endogenous gene, however, R2 and R5 are located ~30 Kb apart from each other, while in the minigene construct the distance between them is reduced to ~2 Kb. Hence, we used AONs complementary to R2 and R5 (AON2-1 and AON2-2, respectively) to confirm that the ultra-long-range base pairing between R2 and R5 also modulates alternative splicing in the endogenous *Ate1* transcript (Figure 3D). RT-PCR and qRT-PCR analyses revealed that 5 nM or a higher concentration of AON2-1 was sufficient to decrease exon 7a usage and increase exon 7b usage in comparison with non-treated cells and cells treated with the control AON, while the proportion of transcripts with double exons remained low (Figure 3E and 3F). The efficacy of AON2-2 was lower since the inclusion of exon 7a decreased to a comparable extent only at higher concentrations of AON2-2 (100 nM). Since the hybridization strengths of AON2-1 and AON2-2 are comparable, the difference in their efficacy could be due to their exposure to R2 and R5 at different times during transcription, i.e., AON2-1 has more time to find its target as R2 is transcribed earlier, while AON2-2 has to compete with R2 to hybridize with R5.

Nonetheless, the effects of AON2-1 and AON2-2 on splicing are concordant with each other and consistent with the results of the mutagenesis. Taken together, they indicate that the RNA structure formed by R2 and R5 is functionally distinct from that of R1R3 and R3R4, and it serves to control the isoform ratio rather than the mutually exclusive choice of exons 7a and 7b.

### Crosstalk between competing and ultra-long-range RNA structures

We have demonstrated that the secondary structure of *Ate1* pre-mRNA contains two distinct modules, R1R3/R3R4 and R2R5, where the former ensures mutually exclusive exon inclusion, and the latter regulates the respective isoform ratio. To investigate how these modules interact with each other, we examined the response of alternative splicing in mutants with disruptive and compensatory mutations within R1, R3, and R4 to the treatment with AON2-1, which blocks the interaction between R2 and R5.

Towards this end, we treated the minigenes carrying mutations in R1, R3, and R4 with AON2-1 and measured the changes in alternative splicing with respect to cells treated with control AON (Figure 4A) and non-treated cells (Figure S2). The effect of AON2-1 was equivalent to that of the point mutations that disrupted the interaction between R2 and R5 (Figure 3A) regardless of mutations changing base pairings within R1R3/R3R4, i.e., it suppressed the inclusion of exon 7a and promoted the inclusion of exon exon 7b without affecting the proportion of double exons (Figure 4A). This experiment demonstrated that the ultra-long-range RNA structure formed by R2 and R5 plays a dominant role in choosing between exons 7a and 7b, while not being directly responsible for their mutual exclusivity.

**Figure 4:**
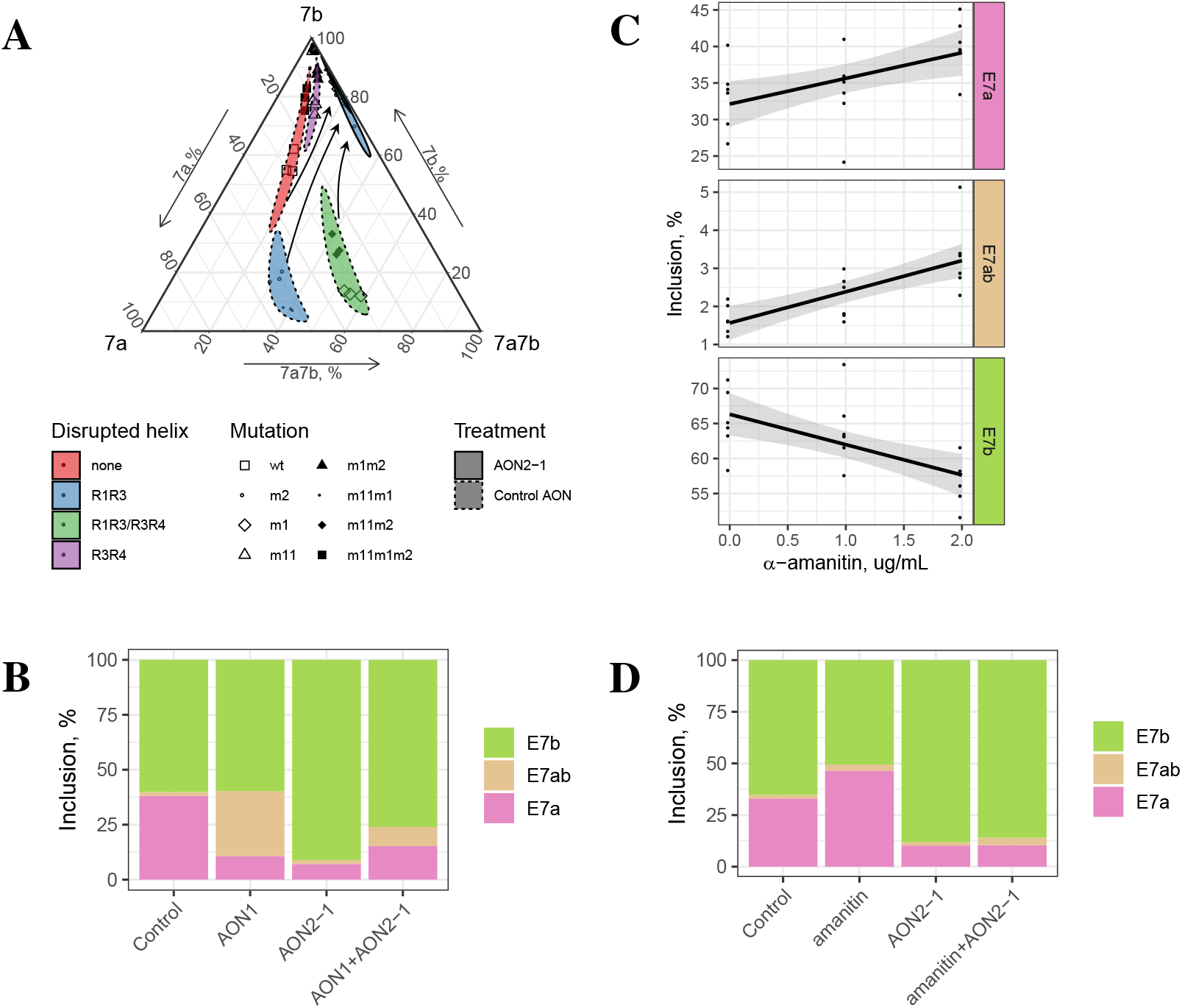
**(A)** Treatment with AON2-1 suppresses the inclusion of exon 7a and promotes the inclusion of exon 7b without affecting the proportion of double exons regardless of mutations in R1, R3, and R4 (Figure 2B). **(B)** The disruption of R1R3/R3R4 base pairing in the endogenous *ATE1* with AON1 increases the proportion of double exons without affecting exon 7b. The disruption of R2R5 with AON2-1 changes exon 7a/7b ratio without inducing double exons. Simultaneous disruption of R1R3, R3R4, and R2R5 yields an additive result. The proportions are averages of two bioreplicates. **(C)** Exon inclusion in response to *α*-amanitin treatment. The scales of y-axes are different between panels. **(D)** Changes of exon inclusion in response to the treatment with AON2-1 and *α*-amanitin. The addition of *α*-amanitin affects exon 7a/7b ratio only when R2R5 base pairing is present.

To discern the interplay between R1R3/R3R4 and R2R5 in the endogenous *Ate1* transcript, we examined the effect of simultaneous disruption of base pairings with AON1 and AON2-1 on exon selection. We treated A549 cells with the combination of 25 nM AON1 and 25 nM AON2-1 and compared the splicing pattern with the effect from single AON treatment and with that of the control AON (Figure 4B). Remarkably, the treatment with AON1 alone increased the proportion of double exons by 28% without changing the rate of exon 7b inclusion, while the treatment with AON2-1 alone conversely increased the inclusion of exon 7b by 31% without introducing double exons. Simultaneous addition of AON1 and AON2-1 led to an intermediate result, in which exon 7b inclusion increased by 16% and the inclusion of double exons increased by 7%. This effect was similar to the response of m1 mutant to the treatment with AON2-1, in which the interaction between R1, R3, and R4 was disrupted by point mutations (Figure 3C).

These results support our hypotheses about functional distinction between two RNA structure modules in *Ate1* pre-mRNA. The module of competing base pairings (R1R3/R3R4) is responsible for mutual exclusivity of exons 7a and 7b, whereas the module of ultra-long-range base pairings (R2R5) controls the isoform balance. Since the latter differs across human tissues (Figure S1A) and disease conditions (Figure 1C), we next questioned which intrinsic factors may modulate the isoform ratio in *Ate1* together with RNA structure.

### Splicing pattern of *Ate1* depends on RNAPII elongation rate

The transcription elongation speed strongly impacts alternative splicing (*39*). It is currently accepted that slow transcription elongation opens a window of opportunity for the upstream splice sites to be recognized, which promotes the inclusion of exons that are otherwise skipped, although in some cases the effect can be the opposite (*40–42*). In our experiments, the disruption of the same RNA structure by AONs blocking R2 and R5 led to different outcomes, which could be a result of co-transcriptional AON binding. We therefore evaluated how the transcription elongation speed influences *Ate1* splicing using *α*-amanitin, a selective inhibitor that interacts with the core subunit of RNAPII and switches transcription to the “slow mode” (*43,44*).

To assess the efficiency of elongation inhibition by *α*-amanitin, we used 28S rRNA/GAPDH ratio measured with qRT-PCR analysis (Figure S3). In the endogenous *Ate1* transcript, the exposure to 2 ug/mL *α*-amanitin led to a notable decrease of exon 7b usage, increased exon 7a usage, and slight elevation of inclusion of double exons (Figure 4C). This pattern was opposite to the effect of AON2-1 treatment, in which the exon 7a/7b ratio has decreased (Figure 3F). In the minigene construct, however, the ratio of exon 7a/7b isoforms did not change significantly (Figure S4). This suggests that transcription elongation slowdown could promote the interaction of R2 with R5 by allowing sufficient time for RNA to fold, and that the absence of the effect in the minigene construct could be related to the shortening of the loop between R2 and R5.

To dissect the interplay between transcription elongation speed and RNA structure, we tested the effect of *α*-amanitin on the endogenous *Ate1* transcript in cells treated with AON2-1 at the concentration disrupting R2R5 base pairing. The addition of AON2-1 abolished the effect of *α*-amanitin, i.e., the inclusion rate of exon 7a decreased, the inclusion rate of exon 7b increased, and double exons were not affected by *α*-amanitin (Figure 4D). This indicates that the increase of exon 7a/7b ratio after *α*-amanitin treatment in the endogenous *Ate1* with intact R2R5 structure was not due to a longer opportunity window for the spliceosome to recognize exon 7a, but rather to a longer time for the RNA structure to fold.

We therefore conclude that the formation of ultra-long-range RNA pairing between R2 and R5 depends on transcript elongation speed, and that the impact from its slowdown on *Ate1* splicing is mediated by the ultra-long-range RNA base pairing R2R5.

## Discussion

Mutually exclusive splicing is among the top five most abundant alternative splicing classes after exon skipping, alternative 5’- and 3’-splice site usage, and retained introns (*45*). Instances of MXEs have been described in diverse phyla including *C. elegans*, *D. melanogaster*, and plants (*28,29,46,47*). Pre-mRNAs of many essential human genes such as glutamate receptor subunits 1-4 (GluR1-4) and voltage-gated sodium channels (SCN genes) undergo mutually exclusive splicing (*48, 49*). MXE clusters often have tissue- and developmental stage-specific expression (*28*), and mutations in them have been linked to hereditary diseases (*50–53*) and cancer (*54, 55*).

Mutually exclusive splicing can be regulated by several distinct mechanisms including spliceosome incompatibility (*56*), steric hindrance of splice sites (*47*), or frame shifting coupled with degradation by nonsense-mediated decay (*57*), but competing RNA secondary structure represents the major mechanism reported in most documented cases (*29*). Here, we demonstrate for the first time an example of MXE in a human gene with two independent structural modules, which have distinct functions: a competing RNA structure module that controls mutually exclusive splicing, and an ultra-long-range base pairing module that spans over 30 kb and regulates the ratio of transcript isoforms. The latter makes *Ate1* a gene with the longest such RNA structure known to date.

The competing base pairings in R1R3 and R3R4 regulate MXE choice by the hindrance of one of the two 3’-splice sites that mediates exposure of only one of them to the spliceosome. On one hand, the relative abundances of these two structures, and consequently the ratio of exon 7a and 7b transcript isoforms, must be proportionate to the difference of their folding energies. On the other hand, mutations in R1 and R4 affect the polypyrimidine tracts and decrease the recognition of exon 7a and 7b by the spliceosome. In all mutants except m2 and m11, the sequences of R1 and R4 either become obstructed by RNA structure or concordantly decrease their affinity to the spliceosome. Consequently, the ratio of exon 7a and 7b isoforms linearly depends on the difference of R1R3 and R3R4 hybridization energies in all mutants except m2 and m11 (Figure 2F). This agrees with the previous findings, in which the rate of exon inclusion inversely correlated with the thermodynamic stability of the surrounding RNA structure in *CFTR* minigene system (*58*). The incomplete reversal of the exon 7a/ 7b ratio in the triple mutant (m1m2m11) is consistent since the balance between the free energies of R1R3 and R3R4 in it was restored to a different level compared to the WT (Figure 2F).

Extensive evidence indicates that speed of transcription elongation may affect the choice of splice sites by changing timing in which they are presented to the spliceosome (*42, 59, 60*). For instance, a DNA-binding factor CTCF can promote inclusion of weak upstream exons by mediating local RNA polymerase II pausing (*61*). Here, we observed that exon 7a/7b ratio responds to the slowdown of RNA Pol II in the presence of R2R5 long-range base pairing, but fails to do so when the base pairing is disrupted. This indicates that beyond co-transcriptional recognition of splice sites, slow transcription elongation also can affect splicing through RNA structure by opening a longer opportunity window for it to fold. In bacterial organisms, kinetic mechanisms that involve RNA structure represent a commonly used strategy of regulating premature transcription termination, in which an RNA element typically senses specific molecules (*62*). In eukaryotes, the kinetics of RNA folding could be influenced by many other factors including changes in the composition of RNA-binding proteins and environmental signals, thus adding to the already complex and dynamic regulatory landscape of alternative splicing.

MXEs within a cluster often share high similarity at the sequence level indicating that some of them could have emerged through tandem genomic duplications (*26*). Furthermore, this mechanism could also generate competing RNA structures by duplicating one of the two arms of an ancestral stem-loop resulting in two selector sequences that compete for the same docking site (*32*). In the case of exons 7a and 7b in *Ate1*, the duplication likely happened after the radiation of Chordata since a homolog of where exon 7a, but not exon 7b is present in invertebrates. Still, it is not possible to track the origin of R1, R3, and R4 since competing RNA secondary structures are usually not conserved across long evolutionary distances (*28*). Interestingly, the length of the intron downstream of exon 7b is decreasing with increasing the evolutionary distance from human (Figure S5), which suggests that the base pairing between R2 and R5 pairing has evolved after exon duplication to counteract the expansion of the intron downstream of exon 7b.

The fact that the ratio of exon 7a/7b is proportional to the difference of hybridization energies of R1R3 and R3R4 despite the former has a larger loop than the latter indicates that *Ate1* pre-mRNA is a highly folded molecule with many more complementary interactions than we have identified here. Chemical RNA structure probing methods are insufficient to determine long-range base pairings since they can only reveal which bases are single-stranded, but cannot identify the interacting partners (*63–67*). Other rapidly emerging technologies such as RNA *in situ* conformation sequencing (RIC-seq) can be used instead to profile long-range RNA structures (*68–71*). The published RIC-seq data for HeLa cells confirm R1R3 and R3R4 base pairings as well as over 50 other base paired regions in exon 7a/7b MXE cluster with a potential impact on splicing (*68*) (Figure S6). Therefore, the regulation of *Ate1* alternative splicing could, in fact, be much more complex than it is described here.

## Conclusion

We demonstrated that the mutually exclusive choice of exons 7a and 7b in the human *Ate1 gene* is regulated by a complex RNA structure composed of two modules: a competing RNA structure module that ensures that one and only one exon is included in the mature mRNA, and an ultra-long-range RNA structure that determines the proportion of transcripts with exons 7a and 7b. The latter can be adjusted using LNA-based antisense oligonucleotides, which opens a broad spectrum of possibilities for targeting alternative splicing of this gene therapeutically.

## Acknowledgements

M.K. and D.S. were supported by Russian Foundation for Basic Research grant 18-29-13020-MK. T.I. was supported by Russian Foundation for Basic Research grant 19-34-90174. S.K.was supported by Russian Foundation for Basic Research grant 18-29-13020-MK and by Skolkovo Institute of Science Technology Research Grant RF-0000000653.

## Materials and Methods

### Minigene construction and mutagenesis

Whole genomic DNA was isolated using Quick-gDNA MiniPrep kit (Zymo Research). *Ate1* minigene was assembled from three fragments: the first one was inserted into pRK5 vector using restriction-free cloning (protocol from (*72*)), and the next two fragments were inserted into resulting plasmid using NEBuilder®HiFi DNA Assembly Cloning Kit (New England Biolabs). Fragments were amplified with primers (Table S1) and Q5 High-Fidelity DNA polymerase (New England Biolab). Minigene mutagenesis was performed using either protocol from (*73*) or using phosphorylated primers that introduced the desired changes with subsequent ligation using Rapid DNA ligation kit (Thermo Scientific). All primers for mutagenesis are listed in Table S1; all PCR reactions were performed using Phusion High-Fidelity PCR Master Mix (Thermo Scientific). All mutants were verified by sequencing.

### Antisense oligonucleotides

LNA/DNA mixmers were designed based on (*38*). Synthesis of LNA/DNA mixmers was carried out by Syntol JSC (Moscow, Russia). The sequences of used AONs are listed in (Table S3).

### Cell culture, transfections and treatments

Human A549 lung adenocarcinoma cell line was maintained in DMEM/F-12 medium containing 10% fetal bovine serum, 50 U/ml penicillin, and 0.05 mg/ml streptomycin (all products from Thermo Fisher Scientific) at 37°C in 5 % CO_2_.

Minigene plasmids were transfected using Lipofectamin 3000 (Invitrogen) with reverse transfection protocol for 24 hours followed by cells harvesting by trypsin-EDTA (Thermo Fisher Scientific). AONs treatment was performed using Lipofectamine RNAiMAX (Invitrogen) on 50-70% confluent cells. It took 48 hours of treatment before cells were harvested. *α*-amanitin (Sigma) was added at concentrations 1 and 2 ug/mL to 50-70% confluent cells. After 24 hours of treatment, cells were harvested.

In experiments when cells were transfected with minigenes and AONs together, conditions were as follows. Plasmids and AONs were mixed together prior to transfection, and then these mixtures were transfected using Lipofectamin 3000 (Invitrogen) to 50-70% confluent cells. After 24 hours of treatment cells were harvested.

Experiments with *α*-amanitin and AONs/minigenes treatments were performed next way. Cells were transfected with AONs/minigenes using reverse transfection. After 12–14 hours of transfection media was changed and *α*-amanitin was added. After 24 hours of *α*-amanitin treatment cells were harvested.

### RT-PCR experiments

Total RNA was extracted using PureLink RNA mini kit (Invitrogen) and treated with RNase-Free DNase I (Thermo Scientific) at 37°C for 60 min with following inactivation at 75°C for 10 min. First-strand cDNA was synthesized using Maxima First Strand cDNA Synthesis Kit for RT-qPCR (Thermo Scientific) according to the manufacturer’s instructions. All primers for the following PCR analysis are listed in (Table S2).

We used competitive RT-PCR analysis for the assessment of the ratio between different splice isoforms in one PCR tube. For each PCR reaction, 20-30 ng of cDNA was used. Reactions were carried out using PCR MM (Thermo Scientific). RT-PCR was carried out under the following conditions: denaturing at 95°C for 3 min, 35 cycles of denaturing at 95°C for 30 sec, annealing at 54°C for 30 sec, and extension at 72°C for 1 min, followed by extension at 72°C for 5 min. The resulting products were analyzed on a 3% agarose gel stained with ethidium bromide and visualized using ChemiDoc XRS+ (Bio-Rad). The proportion of each splice isoform was quantified using the ImageJ software.

For RT-qPCR experiments, we used 20-30 ng of cDNA and the reaction mixture with SYBR Green nucleic acid staining (Invitrogen) on a CFX96 of CFX384 Real-Time systems (Bio-Rad). Minus-RT controls without RT enzyme in the cDNA synthesis reaction were performed in every RT-qPCR analysis. For each mutant and/or treatment experiments, at least 2 biological replicates in PCR triplicates were analyzed. The PCR cycle parameters were as follows: 95°C for 10 min and 35 cycles with denaturation at 95°C for 10 s, annealing at 54°C for 20 s and extension at 72°C for 30 s. For each pair of primers in RT-qPCR analysis, the primer efficiencies were estimated using a calibration curve. The expression of isoforms was calculated by the efficiency method (*74*) with primers efficiencies more than 90% in all cases. The expression levels of all isoforms were normalized to the expression of the constitutive exon in the corresponding sample. Then, the sum of all isoform expression levels was taken as 100% to enable comparative analysis of different biological replicates. We additionally checked the sum of expression levels of all isoforms, and it was not statistically different from the expression level of the constitutive exon in all cases.

### Free energy calculation

The hybridization free energies of the WT and mutant sequences of R1–R5 were computed by the RNAup program from Vienna RNA package with the default settings (*75*).

### Percent-Spliced-In (PSI) calculation

RNA-seq data of poly(A)^+^ RNA were downloaded in BAM format from Genotype-Tissue Expression (GTEx) project (*34*) and The Cancer Genome Atlas (*35*) (8,555 samples from v7 release of GTEx; 731 pairs of samples from TCGA). RNA-seq experiments were processed by IPSA pipeline to obtain split read counts supporting splice junctions (*76*). Split read counts were filtered by the entropy content of the offset distribution, annotation status and canonical GT/AG dinucleotides at splice sites. The exon inclusion rate (Ψ, PSI, or Percent-Spliced-In) was calculated according to the equation

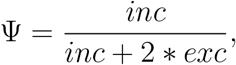

where *inc* is the number of reads supporting exon inclusion and *exc* is the number of reads supporting exon exclusion. Ψ values with the denominator below 20 were considered unreliable and discarded.

### Statistical analysis

The data were analyzed and visualized using R statistics software version 3.4.1 and ggplot2 package (*77*). Ternary diagrams were constructed using the ggtern package (*37*). Confidence intervals (regions) in ternary diagrams were constructed using the Mahalanobis distance (*36*). The statistical significance of log fold change of 7a/7b isoform ratio in TCGA samples was assessed by the Wilcoxon signed rank test with Bonferoni-Holm correction for multiple testing (*FWER* < 0.05). Error bars in the plots and numbers after the ± sign represent 95% confidence intervals. One-sided P-values are reported throughout the paper.

## References

1. Kashina, A. S. Protein Arginylation: Over 50 Years of Discovery. Methods Mol. Biol. 1337, 1–11 (2015).

2. Balzi, E., Choder, M., Chen, W. N., Varshavsky, A. & Goffeau, A. Cloning and functional analysis of the arginyl-tRNA-protein transferase gene ATE1 of Saccharomyces cerevisiae. J. Biol. Chem. 265, 7464–7471 (1990).

3. Yoo, Y. D. et al. N-terminal arginylation generates a bimodal degron that modulates autophagic proteolysis. Proc. Natl. Acad. Sci. U.S.A. 115, E2716–E2724 (2018).

4. Solomon, V., Lecker, S. H., Goldberg, A. L. & Goldberg, A. L. The N-end rule pathway catalyzes a major fraction of the protein degradation in skeletal muscle. J. Biol. Chem. 273, 25216–25222 (1998).

5. Deka, K., Singh, A., Chakraborty, S., Mukhopadhyay, R. & Saha, S. Protein arginylation regulates cellular stress response by stabilizing HSP70 and HSP40 transcripts. Cell Death Discov 2, 16074 (2016).

6. Lamon, K. D., Vogel, W. H. & Kaji, H. Stress-induced increases in rat brain arginyl-tRNA transferase activity. Brain Res. 190, 285–287 (1980).

7. Bongiovanni, G., Fissolo, S., Barra, H. S. & Hallak, M. E. Posttranslational arginylation of soluble rat brain proteins after whole body hyperthermia. J. Neurosci. Res. 56, 85–92 (1999).

8. Kwon, Y. T. et al. An essential role of N-terminal arginylation in cardiovascular development. Science 297, 96–99 (2002).

9. Kurosaka, S. et al. Arginylation-dependent neural crest cell migration is essential for mouse development. PLoS Genet. 6, e1000878 (2010).

10. Rai, R. et al. Arginyltransferase regulates alpha cardiac actin function, myofibril formation and contractility during heart development. Development 135, 3881–3889 (2008).

11. Tanaka, Y. & Kaji, H. Incorporation of arginine by soluble extracts of ascites tumor cells and regenerating rat liver. Cancer Res. 34, 2204–2208 (1974).

12. Chakraborty, G. & Ingoglia, N. A. N-terminal arginylation and ubiquitin-mediated proteolysis in nerve regeneration. Brain Res. Bull. 30, 439–445 (1993).

13. Wang, Y. M. & Ingoglia, N. A. N-terminal arginylation of sciatic nerve and brain proteins following injury. Neurochem. Res. 22, 1453–1459 (1997).

14. Kaji, H. & Kaji, A. Correlated Measurement of Endogenous ATE1 Activity on Native Acceptor Proteins in Tissues and Cultured Cells to Detect Cellular Aging. Methods Mol. Biol. 1337, 39–48 (2015).

15. Lamon, K. D. & Kaji, H. Arginyl-tRNA transferase activity as a marker of cellular aging in peripheral rat tissues. Exp. Gerontol. 15, 53–64 (1980).

16. Rai, R. et al. Arginyltransferase suppresses cell tumorigenic potential and inversely correlates with metastases in human cancers. Oncogene 35, 4058–4068 (2016).

17. Birnbaum, M. D. et al. Reduced Arginyltransferase 1 is a driver and a potential prognostic indicator of prostate cancer metastasis. Oncogene 38, 838–851 (2019).

18. Galiano, M. R., Goitea, V. E. & Hallak, M. E. Post-translational protein arginylation in the normal nervous system and in neurodegeneration. J. Neurochem. 138, 506–517 (2016).

19. Leu, N. A., Kurosaka, S. & Kashina, A. Conditional Tek promoter-driven deletion of arginyl-transferase in the germ line causes defects in gametogenesis and early embryonic lethality in mice. PLoS ONE 4, e7734 (2009).

20. Brower, C. S. & Varshavsky, A. Ablation of arginylation in the mouse N-end rule pathway: loss of fat, higher metabolic rate, damaged spermatogenesis, and neurological perturbations. PLoS ONE 4, e7757 (2009).

21. Kwon, Y. T., Kashina, A. S. & Varshavsky, A. Alternative splicing results in differential expression, activity, and localization of the two forms of arginyl-tRNA-protein transferase, a component of the N-end rule pathway. Mol. Cell. Biol. 19, 182–193 (1999).

22. Rai, R. & Kashina, A. Identification of mammalian arginyltransferases that modify a specific subset of protein substrates. Proc. Natl. Acad. Sci. U.S.A. 102, 10123–10128 (2005).

23. Hu, R. G. et al. Arginyltransferase, its specificity, putative substrates, bidirectional promoter, and splicing-derived isoforms. J. Biol. Chem. 281, 32559–32573 (2006).

24. Schmid, R. et al. The splicing landscape is globally reprogrammed during male meiosis. Nucleic Acids Res. 41, 10170–10184 (2013).

25. Brower, C. S. et al. Liat1, an arginyltransferase-binding protein whose evolution among primates involved changes in the numbers of its 10-residue repeats. Proc. Natl. Acad. Sci. U.S.A. 111, E4936–4945 (2014).

26. Kondrashov, F. A. & Koonin, E. V. Origin of alternative splicing by tandem exon duplication. Hum. Mol. Genet. 10, 2661–2669 (2001).

27. Hatje, K. & Kollmar, M. Expansion of the mutually exclusive spliced exome in Drosophila. Nat Commun 4, 2460 (2013).

28. Hatje, K. et al. The landscape of human mutually exclusive splicing. Mol. Syst. Biol. 13, 959 (2017).

29. Jin, Y., Dong, H., Shi, Y. & Bian, L. Mutually exclusive alternative splicing of pre-mRNAs. Wiley Interdiscip Rev RNA 9, e1468 (2018).

30. Graveley, B. R. Mutually exclusive splicing of the insect Dscam pre-mRNA directed by competing intronic RNA secondary structures. Cell 123, 65–73 (2005).

31. Yue, Y. et al. Long-range RNA pairings contribute to mutually exclusive splicing. RNA 22, 96–110 (2016).

32. Ivanov, T. M. & Pervouchine, D. D. An Evolutionary Mechanism for the Generation of Competing RNA Structures Associated with Mutually Exclusive Exons. Genes (Basel) 9 (2018).

33. Pervouchine, D. D. Towards Long-Range RNA Structure Prediction in Eukaryotic Genes. Genes (Basel) 9 (2018).

34. Melé, M. et al. Human genomics. The human transcriptome across tissues and individuals. Science 348, 660–665 (2015).

35. Weinstein, J. N. et al. The Cancer Genome Atlas Pan-Cancer analysis project. Nat. Genet. 45, 1113–1120 (2013).

36. Reiser, B. Confidence intervals for the mahalanobis distance. Communications in Statistics. Simulation and Computation 30 (2001).

37. Hamilton, N. E. & Ferry, M. ggtern: Ternary diagrams using ggplot2. Journal of Statistical Software, Code Snippets 87, 1–17 (2018).

38. Touznik, A., Maruyama, R., Hosoki, K., Echigoya, Y. & Yokota, T. LNA/DNA mixmer-based antisense oligonucleotides correct alternative splicing of the SMN2 gene and restore SMN protein expression in type 1 SMA fibroblasts. Sci Rep 7, 3672 (2017).

39. de la Mata, M. et al. RNA Polymerase II Elongation at the Crossroads of Transcription and Alternative Splicing. Genet Res Int 2011, 309865 (2011).

40. Saldi, T., Cortazar, M. A., Sheridan, R. M. & Bentley, D. L. Coupling of RNA Polymerase II Transcription Elongation with Pre-mRNA Splicing. J. Mol. Biol. 428, 2623–2635 (2016).

41. de la Mata, M. et al. A slow RNA polymerase II affects alternative splicing in vivo. Mol. Cell 12, 525–532 (2003).

42. Dujardin, G. et al. How slow RNA polymerase II elongation favors alternative exon skipping. Mol. Cell 54, 683–690 (2014).

43. Gong, X. Q., Nedialkov, Y. A. & Burton, Z. F. Alpha-amanitin blocks translocation by human RNA polymerase II. J. Biol. Chem. 279, 27422–27427 (2004).

44. Kaplan, C. D., Larsson, K. M. & Kornberg, R. D. The RNA polymerase II trigger loop functions in substrate selection and is directly targeted by alpha-amanitin. Mol. Cell 30, 547–556 (2008).

45. Wang, E. T. et al. Alternative isoform regulation in human tissue transcriptomes. Nature 456, 470–476 (2008).

46. Yue, Y. et al. Role and convergent evolution of competing RNA secondary structures in mutually exclusive splicing. RNA Biol 14, 1399–1410 (2017).

47. Kuroyanagi, H., Takei, S. & Suzuki, Y. Comprehensive analysis of mutually exclusive alternative splicing in C. elegans. Worm 3, e28459 (2014).

48. Copley, R. R. Evolutionary convergence of alternative splicing in ion channels. Trends Genet. 20, 171–176 (2004).

49. Sommer, B. et al. Flip and flop: a cell-specific functional switch in glutamate-operated channels of the CNS. Science 249, 1580–1585 (1990).

50. Splawski, I. et al. Ca(V)1.2 calcium channel dysfunction causes a multisystem disorder including arrhythmia and autism. Cell 119, 19–31 (2004).

51. Splawski, I. et al. Severe arrhythmia disorder caused by cardiac L-type calcium channel mutations. Proc. Natl. Acad. Sci. U.S.A. 102, 8089–8096 (2005).

52. Bhoj, E. J. et al. Pathologic Variants of the Mitochondrial Phosphate Carrier SLC25A3: Two New Patients and Expansion of the Cardiomyopathy/Skeletal Myopathy Phenotype With and Without Lactic Acidosis. JIMD Rep 19, 59–66 (2015).

53. Mayr, J. A. et al. Deficiency of the mitochondrial phosphate carrier presenting as myopathy and cardiomyopathy in a family with three affected children. Neuromuscul. Disord. 21, 803–808 (2011).

54. David, C. J., Chen, M., Assanah, M., Canoll, P. & Manley, J. L. HnRNP proteins controlled by c-Myc deregulate pyruvate kinase mRNA splicing in cancer. Nature 463, 364–368 (2010).

55. Valletti, A. et al. Identification of tumor-associated cassette exons in human cancer through EST-based computational prediction and experimental validation. Mol. Cancer 9, 230 (2010).

56. Sharp, P. A. & Burge, C. B. Classification of introns: U2-type or U12-type. Cell 91, 875–879 (1997).

57. Jones, R. B. et al. The nonsense-mediated decay pathway and mutually exclusive expression of alternatively spliced FGFR2IIIb and -IIIc mRNAs. J. Biol. Chem. 276, 4158–4167 (2001).

58. Hefferon, T. W., Groman, J. D., Yurk, C. E. & Cutting, G. R. A variable dinucleotide repeat in the CFTR gene contributes to phenotype diversity by forming RNA secondary structures that alter splicing. Proc. Natl. Acad. Sci. U.S.A. 101, 3504–3509 (2004).

59. Ip, J. Y. et al. Global impact of RNA polymerase II elongation inhibition on alternative splicing regulation. Genome Res. 21, 390–401 (2011).

60. Schor, I. E., Gómez Acuña, L. I. & Kornblihtt, A. R. Coupling between transcription and alternative splicing. Cancer Treat. Res. 158, 1–24 (2013).

61. Shukla, S. et al. CTCF-promoted RNA polymerase II pausing links DNA methylation to splicing. Nature 479, 74–79 (2011).

62. Naville, M. & Gautheret, D. Transcription attenuation in bacteria: theme and variations. Brief Funct Genomic Proteomic 8, 482–492 (2009).

63. Ding, Y. et al. In vivo genome-wide profiling of RNA secondary structure reveals novel regulatory features. Nature 505, 696–700 (2014).

64. Rouskin, S., Zubradt, M., Washietl, S., Kellis, M. & Weissman, J. S. Genome-wide probing of RNA structure reveals active unfolding of mRNA structures in vivo. Nature 505, 701–705 (2014).

65. Spitale, R. C. et al. Structural imprints in vivo decode RNA regulatory mechanisms. Nature 519, 486–490 (2015).

66. Flynn, R. A. et al. Transcriptome-wide interrogation of RNA secondary structure in living cells with icSHAPE. Nat Protoc 11, 273–290 (2016).

67. Wan, Y. et al. Landscape and variation of RNA secondary structure across the human transcriptome. Nature 505, 706–709 (2014).

68. Cai, Z. et al. Ric-seq for global in situ profiling of rna–rna spatial interactions. Communications in Statistics. Simulation and Computation (2020).

69. Ramani, V., Qiu, R. & Shendure, J. High-throughput determination of RNA structure by proximity ligation. Nat. Biotechnol. 33, 980–984 (2015).

70. Mortimer, S. A., Kidwell, M. A. & Doudna, J. A. Insights into RNA structure and function from genome-wide studies. Nat. Rev. Genet. 15, 469–479 (2014).

71. Graveley, B. R. RNA Matchmaking: Finding Cellular Pairing Partners. Mol. Cell 63, 186–189 (2016).

72. van den Ent, F. & Löwe, J. RF cloning: a restriction-free method for inserting target genes into plasmids. J. Biochem. Biophys. Methods 67, 67–74 (2006).

73. Edelheit, O., Hanukoglu, A. & Hanukoglu, I. Simple and efficient site-directed mutagenesis using two single-primer reactions in parallel to generate mutants for protein structure-function studies. BMC Biotechnol. 9, 61 (2009).

74. Pfaffl, M. W. A new mathematical model for relative quantification in real-time RT-PCR. Nucleic Acids Res. 29, e45 (2001).

75. Lorenz, R. et al. ViennaRNA Package 2.0. Algorithms Mol Biol 6, 26 (2011).

76. Pervouchine, D. D., Knowles, D. G. & Guigó, R. Intron-centric estimation of alternative splicing from RNA-seq data. Bioinformatics 29, 273–274 (2013).

77. Wickham, H. ggplot2: Elegant Graphics for Data Analysis (Springer-Verlag New York, 2016). URL https://ggplot2.tidyverse.org.

