## Supplementary information for "Multiple competing RNA structures dynamically control alternative splicing in human ATE1 gene"

### Supplementary Figures

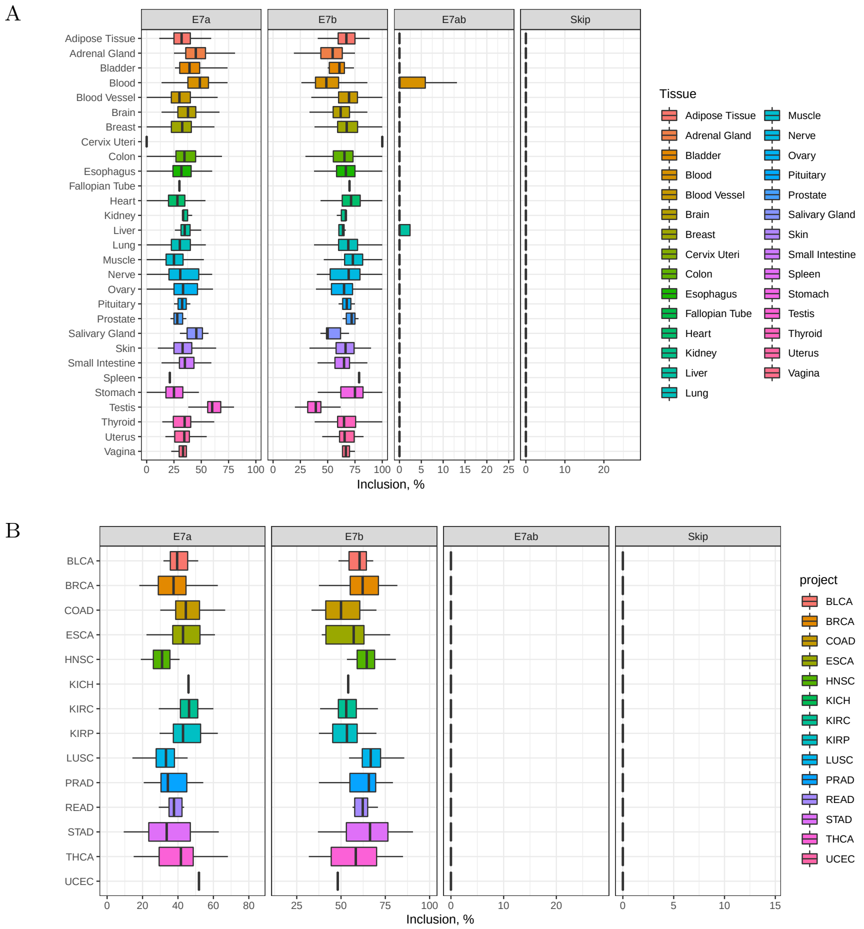

Figure S1: The rate of inclusion of exon 7a, exon 7b, double exons (7a7b), or simultaneous skip of both exons in normal human tissues (A) from Genotype-Tissue Expression (GTEx) project (*1*) and The Cancer Genome Atlas (B) (*2*).

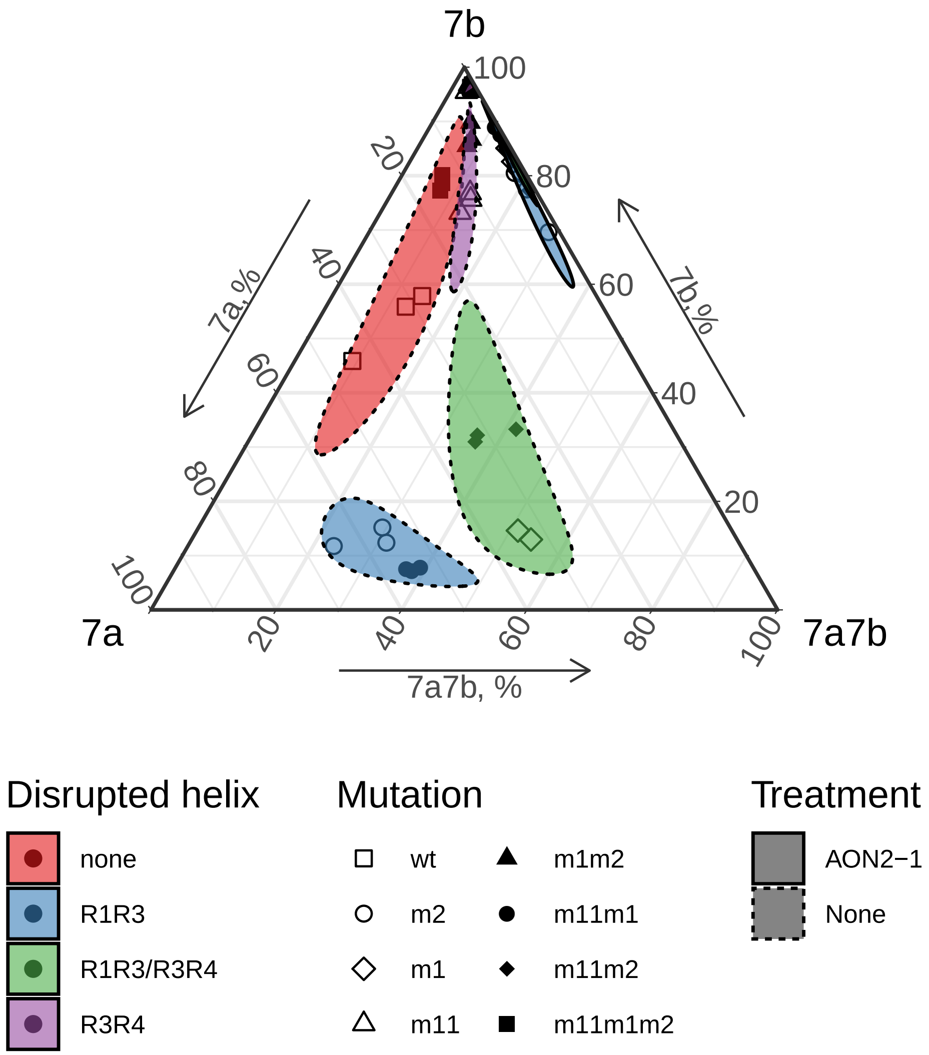

Figure S2: The rate of inclusion of exons 7a, 7b, and double exons (7a7b) in the minigenes carrying mutations in R1, R3, and R4 upon AON2-1 treatment in comparison to non-treated cells. Treatment with AON2-1 suppresses the inclusion of exon 7a and promotes the inclusion of exon 7b without significantly affecting the proportion of double exons.

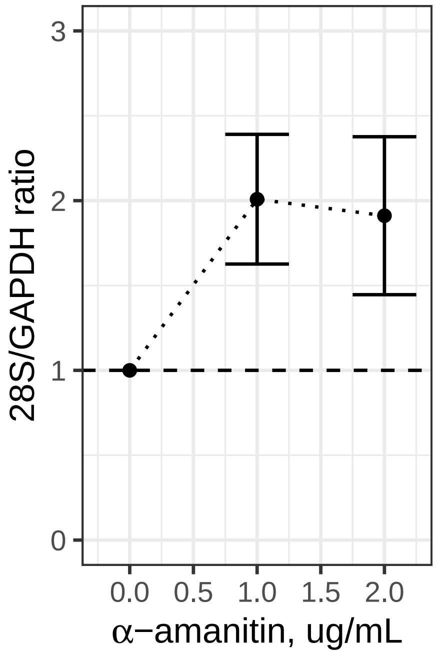

Figure S3: 28S rRNA/GAPDH ratio confirms the efficiency of RNA Pol II elongation inhibition by $\alpha$-amanitin.

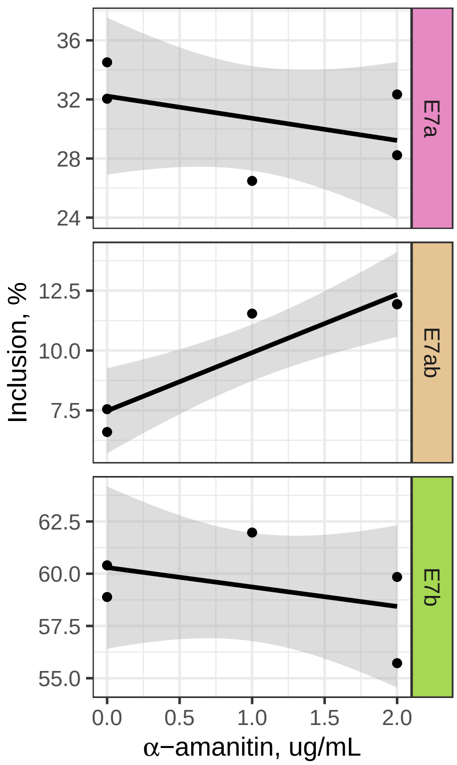

Figure S4: The rate of inclusion of exons 7a, 7b, and double exons (7a7b) in the minigenes did not change significantly upon $\alpha$-amanitin treatment.

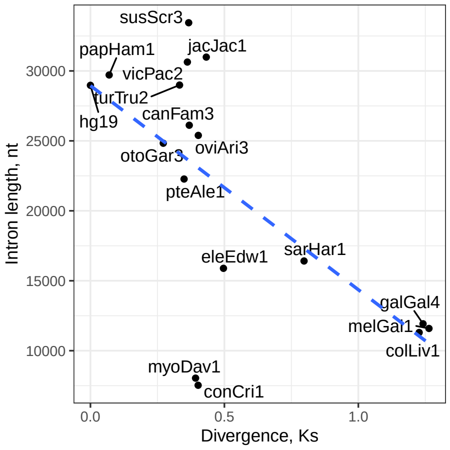

Figure S5: The length of the intron between exon 7b and exon 8 decreases with increasing the evolutionary distance, measured as the average number of substitutions per synonymous site (Ks). The latter was obtained from the phylogenetic tree during the multiz multiple alignment (*3*).

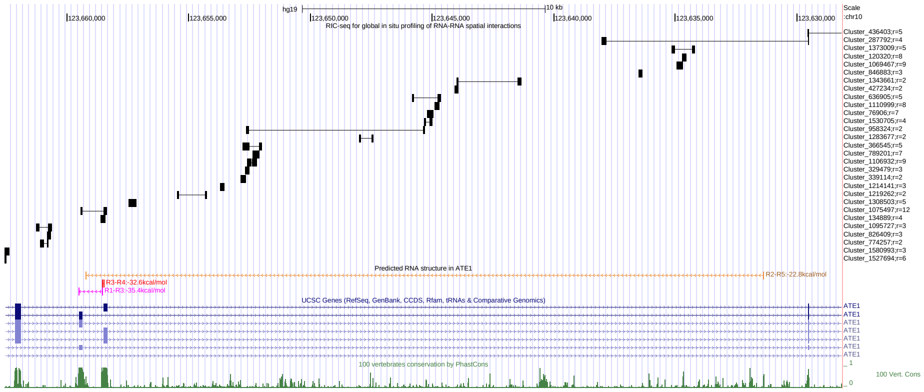

Figure S6: The published RIC-seq data for HeLa cells confirms R1R3 and R3R4 base pairings as well as over 50 other base paired regions in exon 7a/7b MXE cluster with a potential impact on splicing (*4*). The data for mRNAs obtained as a courtesy of Dr. Yuanchao Xue was converted and displayed as a UCSC Genome browser custom track (*5*).

#### Supplementary Tables

| | |
|---|---|
| **ATE1 cloning** | |
| pRK5_Ate1_rf_fwd | GCACCTCGGTTCTATCGATTGAATTCCCCTGAGTAAGCCTCCATGTCGAAAAG |
| pRK5_Ate1_rf_rev | CTGCAGGTCGACTCTAGAGGATCCCCGTATGGCCACTTGATACTTGACATACAA |
| pRK5_ate1_fwd | ATTCCTTTGCGGGGATCCTCTAGAGTCG |
| pRK5_ate1_rev | GGATCCTGGTGTATGGCCACTTGATACTTG |
| ate1_add1_fwd | GTGGCCATACACCAGGATCCACCCGATG |
| ate1_add1_rev | TTCATTGCAGCTGTTGACCTAGAGTGGATGAC |
| ate1_add2_fwd | AGGTCAACAGCTGCAATGAAAGAATCCAAG |
| ate1_add2_rev | GAGGATCCCCGCAAAGGAATCTTGTGAAC |
| **ATE1 mutations** | |
| ate1_m1_fwd | CTCTGATGATATGCTACTCAAGGTGAGGCTACAG |
| ate1_m1_rev | CTGTAGCCTCACCTTGAGTAGCATATCATCAGAG |
| ate1_m2_fwd | ACCACCCTCACCTTGAGTAGCAGTTAAAAAAACAAGT |
| ate1_m2_rev | ACTTGTTTTTTTAACTGCTACTCAAGGTGAGGGTGGT |
| ate1_m11_fwd | ACTCAAGGTGAGGTTAGTACCTGTC |
| ate1_m11_rev | GCAAGGAAAATACAAGCAAATGC |
| ate1_m1_rear_fwd | GGAAGTCTTCGTTATCATCAGAGATGTTTCAATAGCATTTGC |
| ate1_m1_rear_rev | TGAGGCTACAGTAAGATCTCCACAAATC |
| ate1_m4-3_fwd | GACAAAGTTATCTTAAGAGTTAATGTAATGGAACT |
| ate1_m4-3_rev | AAAGTGCAAAATATACCAAAATTTTAAGT |
| ate1_m3-3_fwd | TTGTCAAAGTGCAAATTTTACCTAGTGCC |
| ate1_m3_rev | TGTCCATTAATTCAGTCTTAGG |

Table S1: Cloning and mutagenesis primers

**Minigene**

| | | |
|---|---|---|
| constitutive fragment | pRK5_seq_fwd<br>ate1_rt_rev | CACTCCCAGGTCCAACTG<br>AGCTGGGTTCTGCTGCATTAG |
| isoform with exon 7a | ate1_rt_fwd1<br>ate1_rt_rev1 | TACCAGAGAATGCATCACACA<br>AGGAATCTTGTGAACTGGCTTT |
| isoform with exon 7b | ate1_rt_f2_2<br>ate1_rt_rev2 | TCACACAAGTTAGAGGTGAGGTTAG<br>AGGATCCCCGCAAAGGAATCT |
| isoform with both exons | ate1_ex1_fwd<br>ate1_ex2_rev | ACGTTACCAGATGGTTATTCACAAG<br>GTATGGCCACTTGATACTTGACATA |

**Endogenous ATE1 mRNA**

| | | |
|---|---|---|
| constitutive exon | ate1_const_fwd<br>ate1_const_rev | GCTCTGGTGAACCGTCACATTCAG<br>GACATGGAGGCTTACTCAAATCAGC |
| isoform with exon 7a | ate1_rt_fwd1<br>ate1_rt_rev1 | TACCAGAGAATGCATCACACA<br>AGGAATCTTGTGAACTGGCTTT |
| isoform with exon 7b | ate1_rt_f2_2<br>ate1_rt_r2_c2 | TCACACAAGTTAGAGGTGAGGTTAG<br>CCAAGGGTGAACTGCAAAGGA |
| isoform with both exons | ate1_ex1_fwd<br>ate1_ex2_rev | ACGTTACCAGATGGTTATTCACAAG<br>GTATGGCCACTTGATACTTGACATA |

**RT-PCR primers**

| | |
|---|---|
| ate1_rt_fwd | CCAACCAGCCAAAATCACTCG |
| ate1_ex1_rev | GAACTGCGAACTTGGTGGAGATG |
| ate1_ex2_rev | GTATGGCCACTTGATACTTGACATA |

**primers for 28S/GAPDH qRT-PCR**

| | |
|---|---|
| 28S_fwd | GGAATGCAGCCCAAAGCGG |
| 28S_rev | GGACCCCACCCGTTTACCTCTT |
| Gapdh_fwd | GTCTCCTCTGACTTCAACAGCG |
| Gapdh_rev | ACCACCCTGTTGCTGTAGCCAA |

Table S2: RT-PCR primers

| | |
|---|---|
| ate1_ns | +T*G*+G*A*+A*G*+T*C*+T*T*+C*G*+T |
| ate1_2-1 | +T*G*+C*A*+C*T*+T*T*+C*A*+G*A*+A |
| ate1_2-2 | +A*C*+T*T*+T*C*+T*G*+A*A*+A*G*+T*G*+C*A*+A*A |
| ate1_1 | +T*G*+C*T*+T*C*+T*G*+A*A*+G*G*+T |

Table S3: LNA sequences. DNA base: G,A,T,C. LNA base (red): +G, +A, +T, +C. Phosphorothioated DNA base: G*, A*, T*, C*. Phosphorothioated LNA base (red): +G*, +A*, +T*, +C*.

### References

1. M. Melé, *et al.*, *Science* **348**, 660 (2015).

2. J. N. Weinstein, *et al.*, *Nat. Genet.* **45**, 1113 (2013).

3. W. J. Kent, *et al.*, *Genome Res.* **12**, 996 (2002).

4. Z. Cai, *et al.*, *Communications in Statistics. Simulation and Computation* (2020).

5. B. J. Raney, *et al.*, *Bioinformatics* **30**, 1003 (2014).
